# Endogenous Nitroalkene Exploits Dependence on Autophagy-Lysosome Pathway in PARPi-Resistant Triple Negative Breast Cancer

**DOI:** 10.1101/2025.07.04.663201

**Authors:** Lisa Hong, Sanghoon Lee, Leonard Frisbie, Yaoning Zhao, John J. Skoko, Christopher Merkel, Tetsushi Kataura, Viktor I. Korolchuk, Lan Coffman, Adrian V. Lee, Bruce A Freeman, Francisco J Schopfer, Carola A. Neumann

## Abstract

Lack of DNA double-strand break repair efficiency exquisitely sensitizes cancers to poly-ADP ribose polymerase inhibitors (PARPi). Unfortunately, resistance to PARPi poses an insurmountable challenge for patients. Mechanisms that confer insensitivity to PARPi therapy include enhanced DNA damage repair and autophagy. Natural and non-natural unsaturated fatty acid nitroalkene derivatives (NFA) show anticancer actions that sensitize TNBC cells to PARPi and other DNA-damaging treatments. We reveal that nitro-oleic acid (OA-NO_2_) re-sensitizes PARPi-resistant TNBC cells to PARPi. RNA-seq analysis of clinically relevant *mutBRCA1* PARPi-resistant TNBC cell lines exhibited upregulation in autophagy and lysosomal pathways. Bio-orthogonal analysis identified the autophagy regulator SQSTM1/p62 as a novel OA-NO_2_ target, alkylating two redox-sensitive Cys residues of p62 (Cys105 and Cys113). These Cys are essential for p62 regulation of autophagy and mimicked the effects of p62 Cys105 and Cys113Ala mutants and when alkylated by OA-NO_2_ showed impaired p62 oligomerization, degradation, and inhibition of autophagy. Combination treatment of PARPi-resistant TNBC with a PARPi and OA-NO_2_ synergistically inhibited p62-associated autophagy and lysosome function. These data emphasize the clinical potential of OA-NO_2_ for treating PARPi-resistant TNBC patients.

## INTRODUCTION

Triple-negative breast cancer (TNBC) is one of the most aggressive subtypes of breast cancer. There are currently no approved targeted therapies for TNBC patients who carry loss of function mutations in *BRCA1* and *BRCA2* genes who are eligible for treatment with poly-(ADP-ribose) polymerase (PARP) inhibitors (PARPi) ^1^. PARPi are key targeted therapies for germline *mutBRCA* cancers and are particularly effective due to a synthetic lethality, exploiting a genomic vulnerability in DNA single-strand (SSB) and double-strand break (DSB) repair, such as homologous recombination (HR). They target PARP family proteins, which play essential roles in cell survival and genomic integrity ^2^. The two PARPi currently approved for breast cancer patients expressing *mutBRCA* genes are olaparib (Lynparza) and talazoparib (Talzenna) ^3–6^. Clinical trials show that adjuvant PARPi therapy treatment improves treatment efficacy with fewer side effects than chemotherapy ^6^.

Despite showing clinical promise, PARPi resistance limits effective treatment and patient survival. Thus, improved understanding of PARPi resistance mechanisms aids in the development of more effective therapies for TNBC ^7^. Since *mutBRCA* tumors are sensitive to PARPi, most resistance mechanisms arise from alterations in the DDR pathway, such as the restoration of HR ^7^. As a result, there is clinical interest in combining PARPi with existing and novel HR inhibitors. Also, several FDA-approved PARPi induce autophagy in cancer cells ^8–12^, adding autophagy to the list of PARPi resistance mechanisms ^12–15^. Notably, autophagy inhibition not only increases sensitivity to PARPi ^16,17^, it also reverses PARPi resistance in preclinical models ^11^.

Autophagy is a conserved, catabolic process that maintains cellular homeostasis through the intracellular degradation of cytosolic macromolecules and organelles that is upregulated in response to stress stimuli such as nutrient starvation and DNA damage ^18,19^. Basal autophagy prevents the accumulation of damaged or dysfunctional molecules, such as organelles and misfolded proteins, by facilitating degradation and recycling ^20^. Continuous quality control by autophagy prevents proteotoxicity associated with protein aggregate formation^21^. Of the three main types of autophagy, macroautophagy is the most well-studied and best-characterized form of autophagy. Macroautophagy (autophagic flux), hereafter referred to as autophagy, is a highly organized process characterized by several evolutionarily conserved autophagy-related (ATG) genes and kinases ^22,23^ that initiate autophagic cargo sequestration, fusion, and degradation ^21,24,25^. Despite the potential for autophagy inhibition as a therapeutic intervention, chloroquine (CQ) and hydroxychloroquine (HCQ) are the only FDA-approved autophagy inhibitors ^26^. Even an informed clinical usage of these aminoquinolines, which have a very narrow safe dosing range, presents significant toxicological risks due to impairment of ion transport and organelle trafficking, leading to multi-organ myopathies. In particular, cardiovascular effects and electrolyte derangements induced by CQ/HCQ leads to dysrhythmias induced by QRS prolongation, QTc prolongation and other electrophysiological abnormalities, all underlying increased risk for death ^27,28^.

Small molecule nitroalkenes show potential as a treatment for TNBC, both as a standalone therapy and in combination with PARPi and other DNA-directed drug strategies ^29,30^. Electrophilic nitroalkenes react reversibly with hyper-reactive Cys by Michael addition, thus modulating protein structure and function ^31–33^. The NFA 10-nitro-9(*E*)-octadec-9-enoic acid (OA-NO_2_) displays anti-inflammatory properties that are currently being tested in a phase II clinical trials for obesity-related asthma (NCT03762395) ^34,35^. In concert with derisking OA-NO_2_ for clinical administration, we discovered anti-cancer effects of OA-NO_2_ in several preclinical models ^29,30,36^. Both OA-NO_2_ and synthetic non-natural nitroalkenes such as 7- and 8-nitrononadec-7(*E*)-enoic acid inhibit the HR recombinase RAD51 by selectively adducting the C-terminal Cys319. This reaction confers synthetic lethality of NFA when administered in combination with DNA-directed therapies such as PARPi, a standard of care for many cancers^29,30^.

To characterize the mechanisms by which OA-NO_2_ enhances tumor cell killing by PARPi, we examined the effects of OA-NO_2_ on tumor cell PARPi resistance and autophagy. Chemoproteomics strategies revealed another functionally-significant Cys target of OA-NO_2_, p62 (SQSTM1). We also show that, compared to parental or PARPi-sensitive cells, PARPi-resistant TNBC cells exhibit higher levels of basal autophagy, which can be targeted through autophagy inhibition.OA-NO_2_ inhibited autophagy-lysosome functions by alkylating Cys105 and Cys113 of p62. Inhibition of autophagy by OA-NO_2_ also synergized PARPi killing of in PARPi-resistant TNBC cells. These data reveal a novel role for OA-NO_2_ and other small molecule nitroalkenes in targeting lysosomal function, autophagy-mediated chemoresistance, and the resensitization of drug-resistant TNBC cells to PARPi.

## RESULTS

### PARPi induce autophagy in TNBC

PARPi are utilized in the treatment in breast, ovarian, prostate, and pancreatic cancers ^37^, acting in part by inducing autophagy ^8–13,16,38^. MDA-MB-231 TNBC cells, which are *wtBRCA* and HR-proficient, were employed as a model for PARPi resistance, with HR reversion frequently an underlying mechanism. A mouse xenograft model was used to investigate the effects of PARPi and OA-NO_2_ as a combination treatment. Following tumor formation, either vehicle, talazoparib, OA-NO_2_, or a combination of talazoparib and OA-NO_2_ were gavaged daily (**Supplemental Fig. 1a, b**). While the combination treatment significantly reduces tumor growth compared to single-agent therapy alone, talazoparib failed to impede tumor growth (**Fig. 1a**). To next study the effects of PARPi (talazoparib and olaparib) on TNBC cell growth, we analyzed cellular proliferation over time by generating MDA-MB-231 cells stably expressing a nuclear-restricted red fluorescent protein (RFP). After a single treatment with PARPi, there was a significant reduction in cell proliferation rates (**Fig. 1b**). Using olaparib and talazoparib concentrations significantly greater than calculated IC50 values (**Fig. 1c**), these cells continued to proliferate, which correlated with cell volume enlargement and vacuole-like appearances (**Supplemental Fig. 1c**) resembling autophagy-like morphologies ^39,40^.

**Figure 1.**
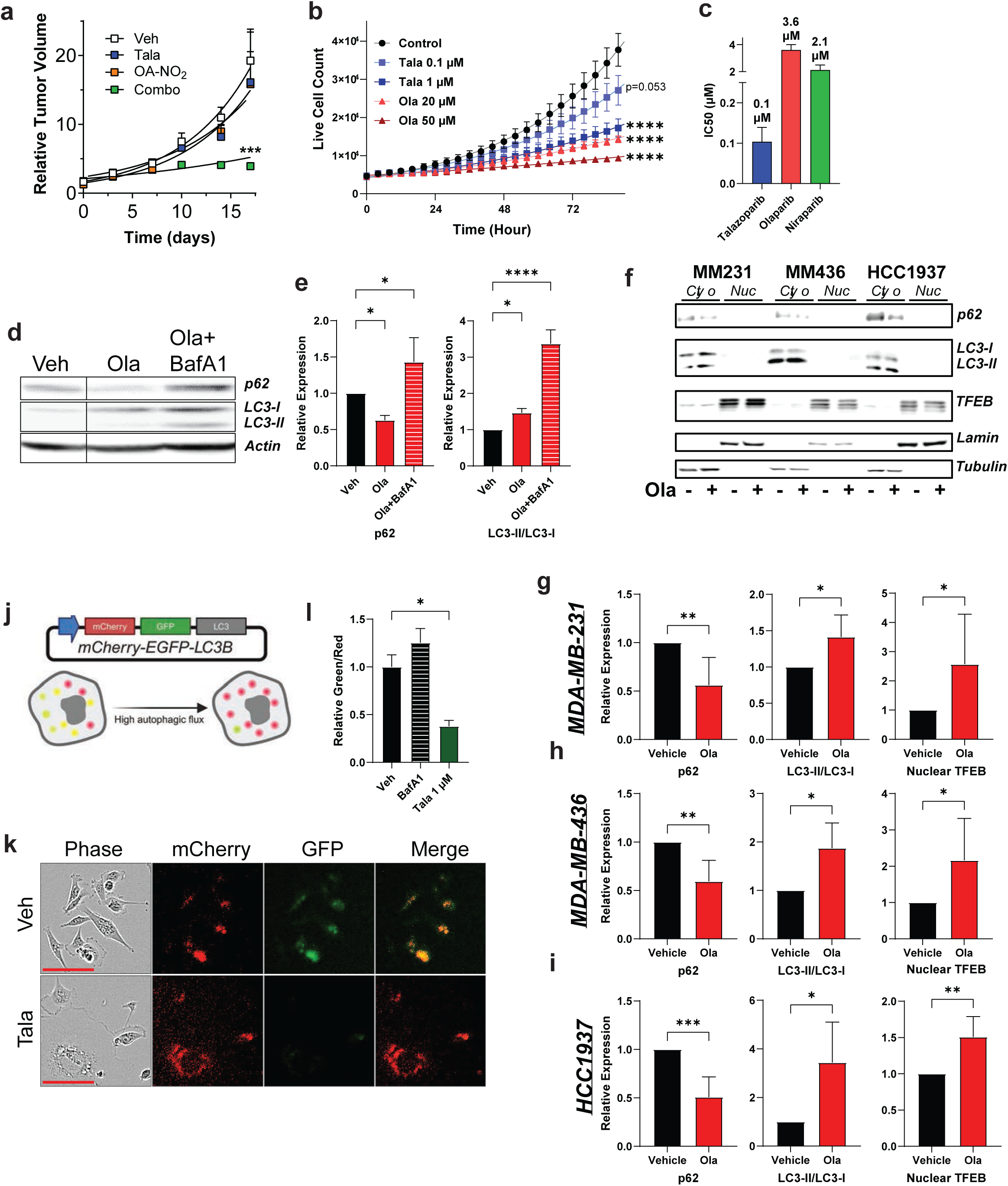
PARPi Induce Autop hagy in TNBC cells.

To examine whether PARPi induce autophagy in TNBC cells, MDA-MB-231 cells were treated with olaparib and then immunoblotted to detect possible changes in autophagy markers. Autophagy inhibition by bafilomycin A1 (BafA1) was used as a control. Olaparib induced p62 protein reduction and further increased LC3-II turnover, indicating autophagy induction (**Fig. 1d, e**). Talazoparib also induced a similar phenotype in MDA-MB-231 (**Supplemental Fig. 1d, e**) and Hs578T (**Supplemental Fig. 1f, g**) TNBC cell lines.

To further establish a potential role for olaparib-induced autophagy in TNBC cells, the BRCA1-mutant TNBC cell lines MDA-MB-436 and HCC1937 ^8,41^ were also analyzed. Olaparib treatment lowered p62 protein levels, increased LC3-II turnover, and increased nuclear protein levels of transcription factor EB (TFEB), a crucial regulatory component in the autophagy-lysosome pathway (**Fig. 1f-i)** ^42^.

A dynamic dual-fluorescent reporter (mCherry-EGFP-LC3B) assay ^43^ expressing the autophagosome marker LC3 and linked to tandem-tagged fluorescent mCherry and GFP proteins was then studied. These cells which constitutively express the pH-stable mCherry show that under conditions of autophagic flux or induced autophagy, the autophagosome fuses with the lysosome, and the acidic environment quenches the GFP signal, resulting in greater red fluorescence. The GFP signal will lead to more yellow foci if the autolysosome fails to form or degrade cargo under autophagy inhibition (**Fig. 1j)** ^43^. Following talazoparib treatment, there was diminished GFP fluorescence compared to the vehicle, indicating the progression of autophagic flux. BafA1 treatment exhibited greater green-to-red fluorescence (**Fig. 1k, l)**. Taken together, these results confirm that PARPi induces autophagy in TNBC cells.

### Characterization of PARPi-resistant TNBC

Overcoming PARPi resistance in TNBC treatment is an unmet clinical need. To closely model the patient population in TNBC approved for PARPi treatment, we generated PARPi-resistance in two TNBC cell lines that carry mutations in the *BRCA1* gene, MDA-MB-436 and HCC1937. Based on previous methods, resistance was conferred through continuous exposure to increasing concentrations of the PARPi olaparib, talazoparib, and niraparib ^44^. HR-proficient MDA-MB-231 cells were included as a control. After gradually increasing the concentration of PARPi over the course of 8 months, PARPi resistance was confirmed by determining the IC50 values for talazoparib, olaparib, and niraparib (**Table 1)**. Cell lines were deemed resistant (PARPiRes) when their PARPi IC50s at least two-fold greater than the IC50s of their respective naïve (parental) cell lines.

Overall, PARPi-resistant cell lines exhibited an enlarged cell volume and increased vacuole formation. The MDA-MB-231 and MDA-MB-436 talazoparib-resistant (TalaRes) and olaparib-resistant (OlaRes) cells also demonstrated a more spindle-like morphology than parental cells (**Supplemental Fig. 2a**). Next, we compared the PARPi IC50 values of parental and PARPi-resistant TNBC cell lines. As shown in **Table 1**, each PARPi-resistant cell line had a greater fold change of IC50 compared to their respective parental cell line, except for MDA-MB-231 niraparib-resistant cells (NiraRes). Notably, the MDA-MB-436 PARPi-resistant cell lines exhibited a fold resistance of over 300 against each of the three PARPi, indicating PARPi cross-resistance (**Fig. 2a**), as previously ^45,46^. Colony formation assays further confirmed this resistance to PARPi (**Fig. 2b and Supplemental Fig. 2b-d**). Olaparib treatment reduced the colony number in parental MDA-MB-436 cells; however, it had a minimal effect on TalaRes and OlaRes colony formation for both cell lines, indicating that these cells proliferate despite PARPi treatment. Lastly, cell cycle analysis also demonstrated that PARPi treatment in PARPi-resistant TNBC did not cause significant arrest in the G2/M phase (**Fig. 2c-e**).

**Figure 2.**
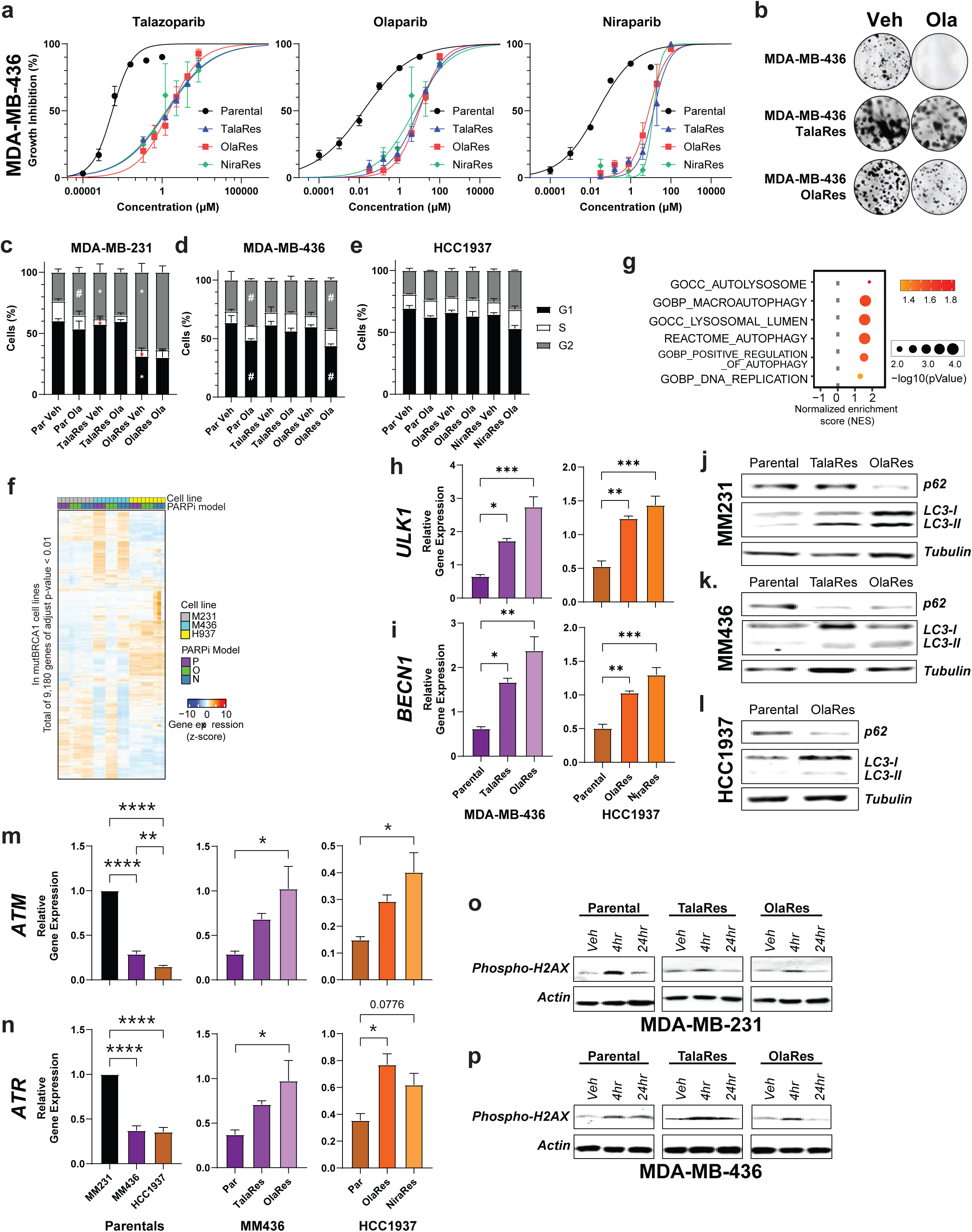
Development of PARPi Resistant TNBC Cell Lines.

RNAseq identified transcriptomic alterations corresponding to drug resistance in PARPi-resistant TNBC cells. From a broad analysis, the gene expression profile heatmap and principal component analysis (PCA) revealed that the individual TNBC cell lines are transcriptomically distinct from one another; however, the MDA-MB-436 TalaRes and OlaRes groups are clustered with *wtBRCA* MDA-MB-231 (**Fig. 2f** and **Supplemental Fig. 2e**). Notably, compared to their respective parental cell lines, the *mutBRCA1* MDA-MB-436 and HCC1937 OlaRes cells, but exclusive in *wtBRCA1* TNBC, shared enriched hallmark pathways, such as K-Ras signaling, inflammatory response, and EMT signaling (**Supplemental Fig. 2f**).

We also identified the enrichment of autophagy-related pathways in PARPi-resistant mutBRCA1 MDA-MB-436 and HCC1937 OlaRes cells against their respective parental cells. There was a positive normalized enrichment score (NES) for lysosomal and autophagy regulatory pathways (**Fig. 2g and Supplemental Fig. 2g**). This finding was validated by RT-qPCR of autophagy-related genes, including *ULK1*, *BECN1*, and *ATG5*, which all were elevated in the PARPi-resistant cells compared to the respective parental cell lines (**Fig. 2h, i** and **Supplemental Fig. 3a-c**). Examining common autophagy-related markers by immunoblotting further confirmed that PARPi resistance induces autophagy as levels of p62 protein were decreased and LC3-II turnover was increased in PARPi-resistant cell lines compared to the parental cell lines (**Fig. 2j-l**).

Since HR restoration is clinically one of the most studied PARPi-resistance mechanisms ^7^, parental and PARPi-resistant counterparts were analyzed for the expression of HR markers. RT-qPCR analysis demonstrated that parental *mutBRCA1* TNBC cells have decreased expression of key DDR and DSB repair genes compared to *wtBRCA* MDA-MB-231 cells (**Fig. 2m, o** and **Supplemental Fig. 3d-f**). Conversely, the PARPi-resistant *mutBRCA1* TNBC cell lines exhibit greater expression of these genes than their respective parental cell lines, suggesting HR restoration in PARPi-resistant *mutBRCA1* cell lines, similar to MDA-MB-231 cells. In support of this, irradiated PARPi-resistant *mutBRCA1* MDA-MB-436 cells showed decreased γH2A.X phosphorylation on Ser136 compared to non-irradiated PARPi-resistant cells, which mirrored the γH2A.X phosphorylation in *wtBRCA* MDA-MB-231 cells (**Fig. 2o, p**).

### OA-NO_2_ targets functionally significant Cys in the autophagy regulator protein p62

To determine the role of OA-NO_2_ in modulating autophagy, MDA-MB-231 cells were treated with OA-NO_2_ +/- the autophagy inhibitor BafA1. Similar to BafA1, OA-NO_2_ increased p62 and LC3-II protein expression (**Fig. 3a, b**). OA-NO_2_ also increased p62 expression in parental and PARPi-resistant MDA-MB-436 and HCC1937 cell lines (**Fig. 3c**). The dual-fluorescent autophagy reporter also demonstrated that OA-NO_2_ phenocopies BafA1 treatment by increasing GFP fluorescence compared to vehicle (**Fig. 3d**), supporting that OA-NO_2_ inhibits autophagic flux. Multiple autophagy-related proteins contain redox-sensitive cysteine residues, with p62/SQSMT1 the best characterized ^47–55^.

**Figure 3.**
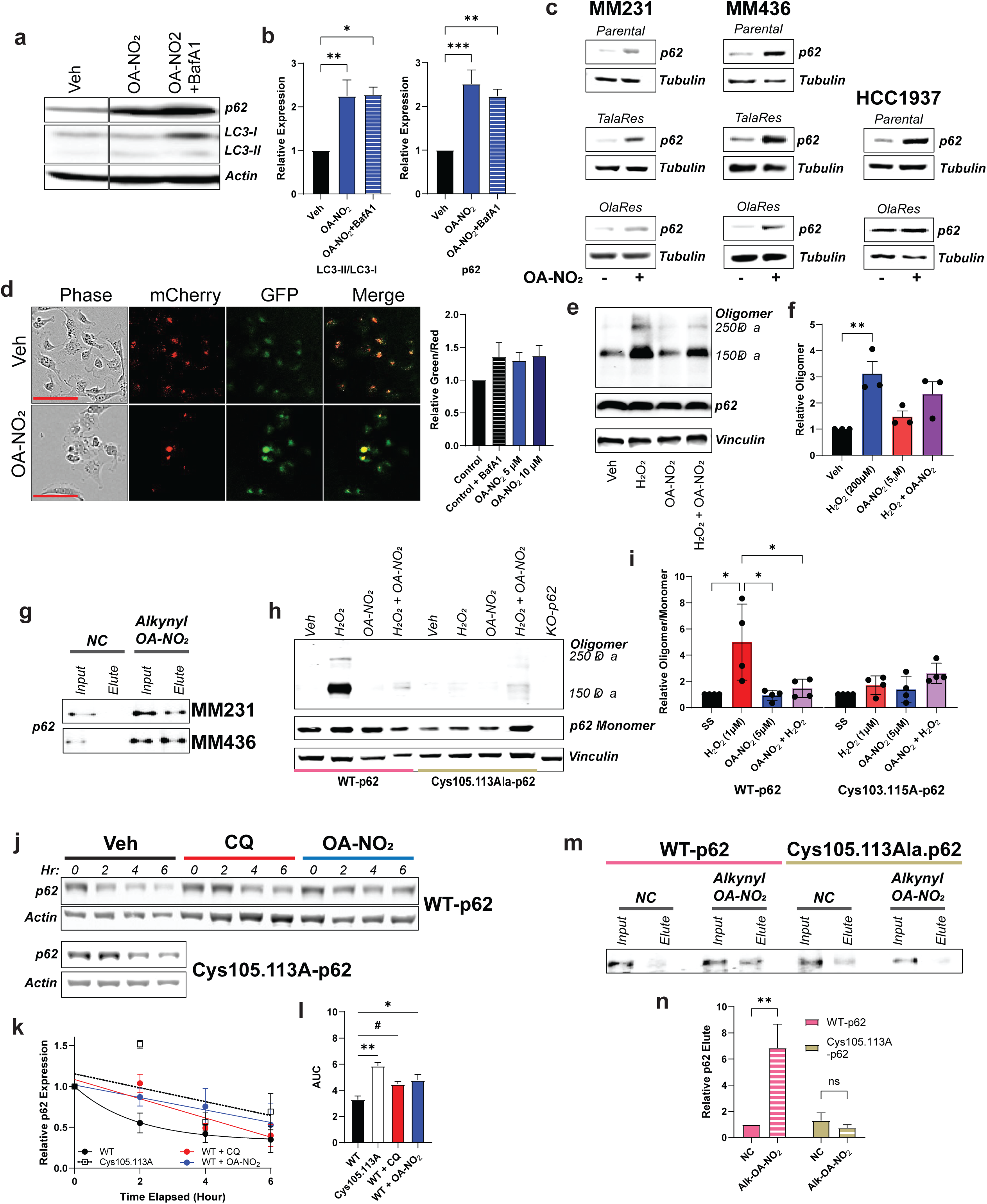
Nitro-oleic acid imp airs autophagic flux by targeting redox-sensitive autophagy proteins.

Adaptor protein p62 activity and protein stability are regulated by oxidative stress through conserved cysteine residues essential for self-oligomerization and protein stability ^48,51,52,56^. We examined if OA-NO_2_ inhibits p62 oligomerization. HEK293T expressing HA-tagged p62 were treated with H_2_O_2_ dosage that induces p62 oligomerization ^48^, which led to p62 dimer and oligomer formation (**Fig. 3e, f**). OA-NO_2_ treatment reduced oligomer formation in H_2_O_2_-treated samples, suggesting that OA-NO_2_ prevents p62 oligomerization by targeting p62 cysteine residues. To examine directly if OA-NO_2_ targets cysteines in p62, we employed a copper-assisted azide-alkynyl-OA-NO_2_ cycloaddition (CuAAC) chemoproteomics approach to enrich alkynyl-OA-NO_2_ targets (**Supplemental Fig. 4a, b**). A variety of click chemistry ^57–59^. CuAAC utilizes copper (I) ions for rapid and efficient bio-orthogonal labeling of proteins under physiological conditions, catalyzing a Huisgen-1,3-dipolar cycloaddition reaction ^60^. Following CuAAC methods ^61–63^, p62 was pulled down and identified to have undergone OA-NO_2_ cysteine alkylation in both MDA-MB-231 and MDA-MB-436 cells (**Fig. 3g)**. Other autophagy-related proteins were also pulled down **(Supplemental Fig. 4c**).

Considering the role of p62 cysteine residues ^47–55,64^ in autophagy p62 protein sequences were compared in 11 organisms ^65^. From this analysis, 14 cysteine residues were evolutionarily conserved (**Table 2**), and more highly conserved in vertebrates (**Supplemental Fig. 4d**). In particular, Cys 105 and 113 have been documented to be crucial for p62 oligomerization and the degradation of substrates during autophagy ^48,56^. The progression from monomer to filament ^66^ promotes p62 autophagosomal cargo linking during autophagy (**Supplemental Fig. 4d**).

To further scrutinize the relationship between OA-NO_2_, p62, and the regulation of autophagy, the function of Cys105 and Cys113 in p62 oligomerization was investigated. Using *p62^-/-^* mouse embryonic fibroblasts (MEF) that express either FLAG-p62WT or FLAG-p62Cys105.113Ala, we induced p62 oligomerization with H_2_O_2_ as previously described ^48^. As expected, Cys105.113Ala mutants failed to exhibit p62 oligomerization in response to oxidative stress (**Fig. 3h, i**). Interestingly, OA-NO_2_ significantly blocked H_2_O_2_-induced oligomerization in WT-p62 expressing MEFs (**Fig. 3h, i**), suggesting that OA-NO_2_ targets Cys105.113 residues. We next evaluated whether OA-NO_2_ treatment prevents p62 degradation. To do so, the MEFs were treated with H_2_O_2_ to induce autophagy before cycloheximide (CHX) treatment. Autophagy inhibition was induced by a combination treatment of H_2_O_2_ and CQ or OA-NO_2_. As expected, p62 degradation was impaired by OA-NO_2_ treatment, which phenocopied Cys105.113Ala MEFs (**Fig. 3j-l**).

Finally, to determine a direct interaction between OA-NO_2_ and these cysteine residues, the click chemistry approach described previously (**Supplemental Fig. 4b**) was utilized in MEFs expressing either FLAG-WTp62 or FLAG-p62Cys105.113Ala. While OA-NO_2_ pulled down p62 in WTp62 MEFs, it was unable to pull down Cys105.113Ala-p62 (**Fig. 3m, n**). These results confirm that OA-NO_2_ reacts with either both or one of the Cys105.113Ala-p62 residues of p62 to regulate p62 activity.

### OA-NO_2_ resensitizes TNBC to PARPi through autophagy inhibition

OA-NO_2_ resensitizes PARPi-resistant TNBC by inhibiting p62 activity. OA-NO_2_ treatment also impaired p62 oligomerization in our parental and PARPi-resistant TNBC cell lines (**Supplemental Fig. 4e-g)**. Consistently, H_2_O_2_-induced p62 dimerization and oligomerization were substantially inhibited by the addition of OA-NO_2_ in all parental and PARPi-resistant cell lines. Also, pathway enrichments were induced by OA-NO_2_ in the PARPi-resistant *mutBRCA1* TNBC cell lines. RNAseq analysis of MDA-MB-436 OlaRes cells revealed decreased expression of DDR and autophagy-related pathways after OA-NO_2_ treatment (**Fig. 4a** and **Supplemental Fig. 5a**). Similar results were observed in the HCC1937 OlaRes cells (**Fig. 4b** and **Supplemental Fig. 5b**). Additionally, OA-NO_2_ did not induce significant cell cycle changes (**Supplemental Fig. 5c**), as previously ^29^.

**Figure 4.**
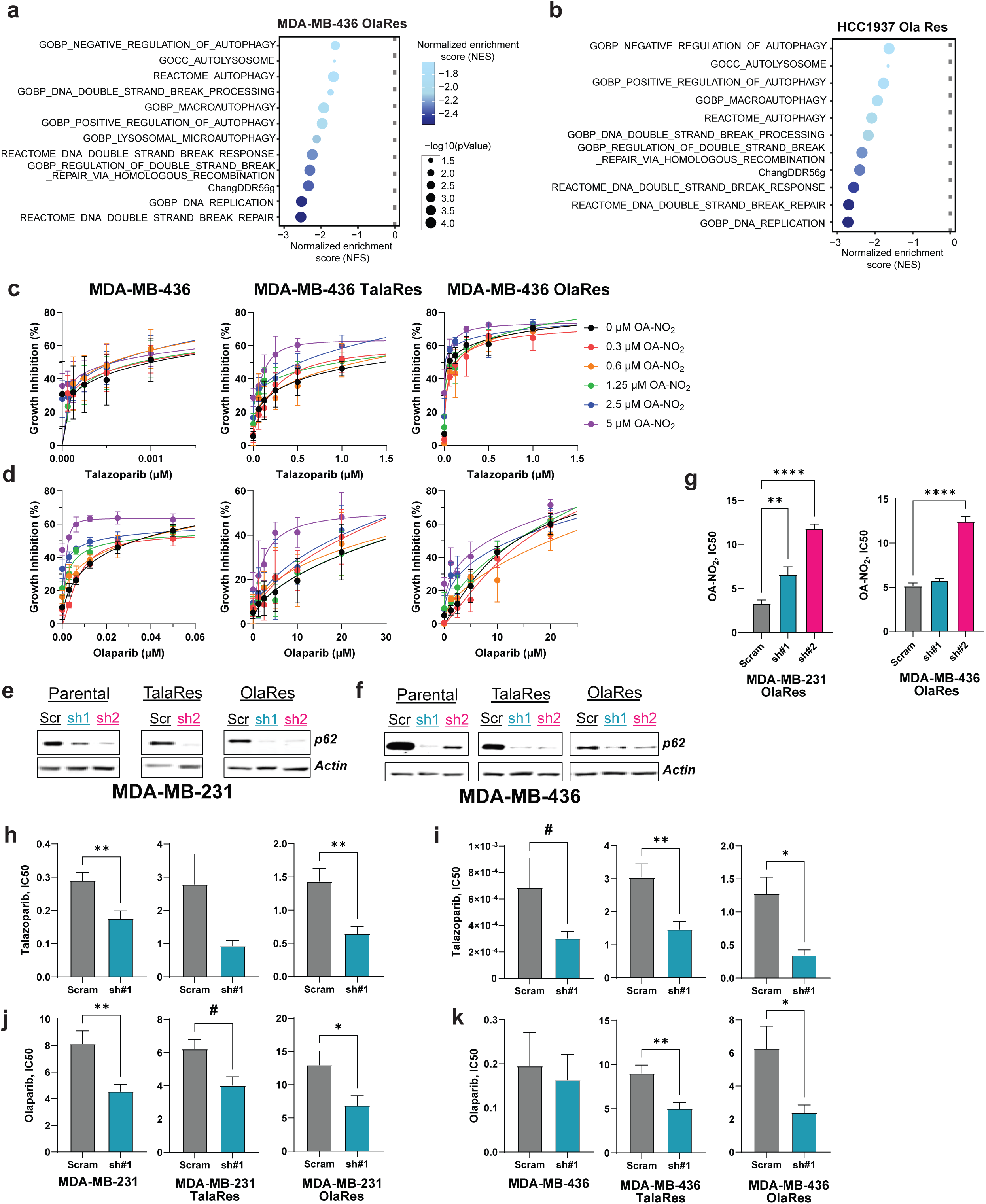
PARPi-Resistant TNBC Resensitized to PARPi through OA-NO_2_ alkylation of p62.

To investigate whether OA-NO_2_ treatment exhibits greater cytotoxicity in PARPi-resistant *mutBRCA1* TNBC compared to parental TNBC cells, combination treatments of PARPi and OA-NO_2_ were conducted while analyzing the resulting highest single agent (HSA) synergy scores. Concentrations for combination indices were determined from PARPi IC50 values (**Table 1**) and OA-NO_2_ IC50 values (**Table 3**).Compared to parental cell lines, greater growth inhibition with talazoparib and olaparib occurred with increasing concentrations of OA-NO_2_ (**Fig. 4c, d**). The results indicate that the combination treatment of PARPi and OA-NO_2_ induces a more pronounced synergistic effect in PARPi-resistant TNBC than in parental TNBC (**Table 4**). This also supports that autophagy inhibition by OA-NO_2_ is a vulnerable target for PARPi resistance. To further validate this, combination indices were determined for PARPi and CQ treatment. Similar synergy scores were found, demonstrating that the combination treatment had a more significant synergistic effect in PARPi-resistant TNBC cells than their respective parental TNBC cell lines (**Table 5**). These results indicate that autophagy is an exploitable pathway to re-sensitize resistant TNBC to PARPi.

To affirm that OA-NO_2_-mediated p62 inhibition contributes to this increased sensitivity, TNBC cell lines (parental and PAPRi-resistant) were generated with decreased p62 expression through shRNA knockdown. It was hypothesized that the knockdown of p62 expression would mimic targeting p62 activity by OA-NO_2_. After the generation of shRNA TNBC cell lines, immunoblot analysis confirmed the knockdown of p62 (**Fig. 4e, f**). A weak correlation was observed between p62 knockdown and TFEB expression (**Supplemental Fig. 5e-g**). Calculated IC50 values for OA-NO_2_ in shRNA p62 knockdown cell lines revealed that p62 knockdown confers resistance to OA-NO_2_ (**Fig. 4g** and **Supplemental Fig. 5d),** reinforcing that p62 is an OA-NO_2_ target. Moreover, the knockdown of p62 conferred increased sensitivity to talazoparib and olaparib in parental and PARPi-resistant TNBC cell lines (**Fig. 4h-k**). This observation aligns with the synergy scores, indicating autophagy inhibition sensitizes PARPi-resistant TNBC to PARPi.

### Lysosomal degradation as a vulnerable pathway in PARPi resistance

During autophagic flux, the lysosome plays a key role by fusing with the autophagosome to form the autolysosome, degrading autophagic substrates. Thus, the completion of the autophagy process is dependent on the lysosome ^67^. The lysosome has been associated with drug resistance in cancer by sequestering hydrophobic weak bases-like chemotherapeutics, resulting in increased numbers of lysosomes ^68^. In TNBC, a recent report attributed this mechanism to CDK4/6 inhibitor resistance ^69^.

Pathway enrichment analysis indicated lysosomal lumen pathway enrichment in the OlaRes *mutBRCA1* TNBC against parental counterparts (**Fig. 2g**). Gene ontology (GO) analysis revealed an upregulation of transcripts related to the lysosomal lumen in both *mutBRCA1* TNBC PARPi-resistant groups (**Fig. 5a** and **Supplemental Fig. 6a**). RT-qPCR also partially validated these results, showing significantly increased mRNA expression of lysosomal-associated membrane protein 2 (LAMP2) (**Fig. 5b**). Similarly, compared to their respective parental cell lines, PARPi-resistant cells exhibit greater expression of nuclear TFEB (**Fig. 5c-e**). These findings suggest that PARPi-resistant cells exhibit increased lysosomal biomass.

**Figure 5.**
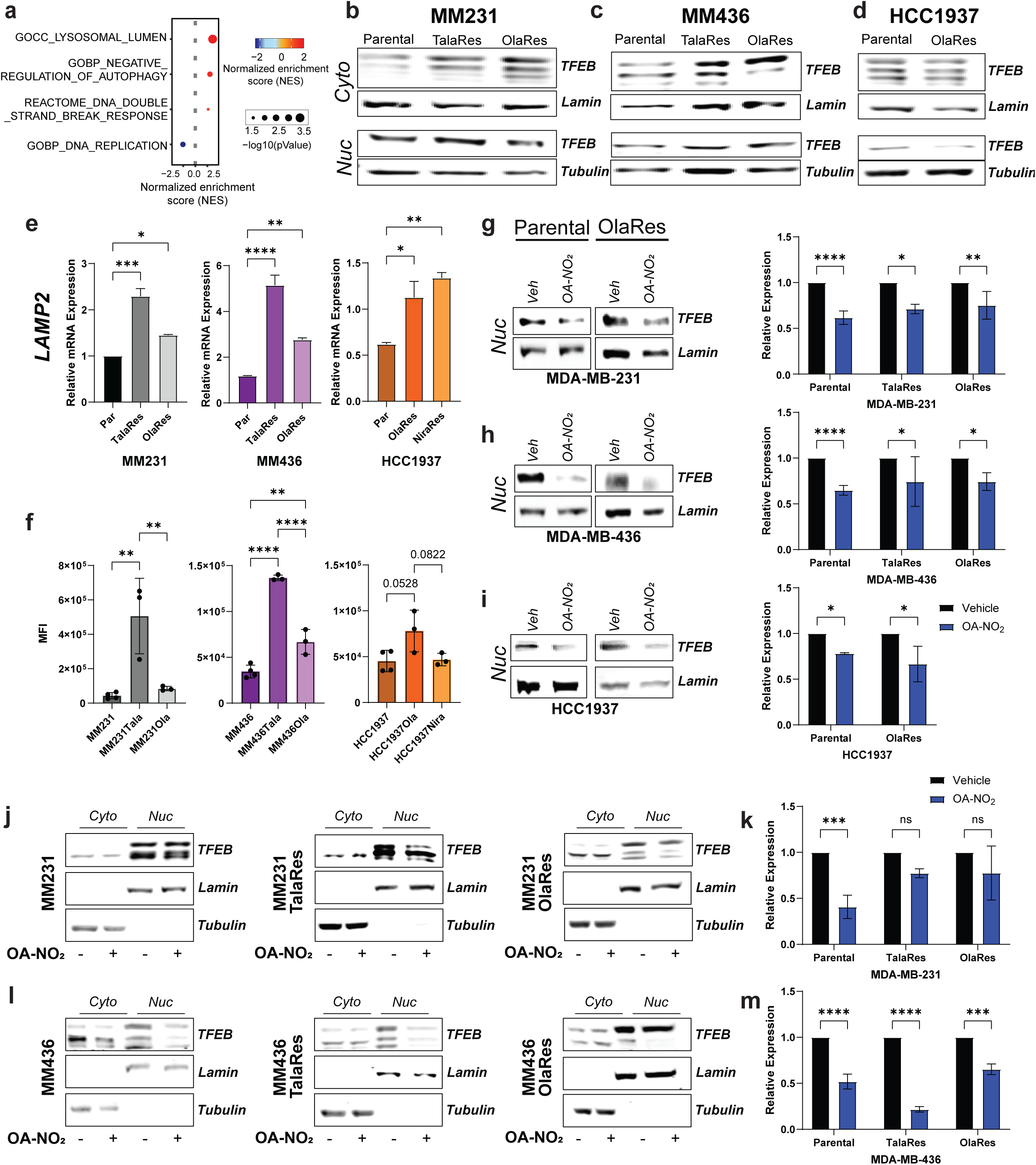
Increased Lysosome Biomass Associated with PARPi Resistance is Targeted by OA-NO_2_.

LysoTracker, a fluorescent dye that stains specifically for lysosomal structures and other acidic organelles revealed a significant increase in the geometric mean fluorescence intensity (MFI) in PARPi-resistant TNBC cells compared to parental controls (**Fig. 5f**). No significant difference was found between parental TNBC cells (**Supplemental Fig. 6b**). This result was further confirmed by decreased CQ and BaFA1 IC50 values in the PARPi-resistant cells compared to the parental cell lines, signifying greater sensitivity to lysosomal inhibition (**Supplemental Fig. 6c-d**). These findings indicate that the PARPi-resistant TNBC cells have a higher lysosomal dependence Both parental and PARPi-resistant TNBC exhibited decreased nuclear TFEB protein expression after OA-NO_2_ treatment (**Fig. 5g-i**). This reduced expression was the result of decreased nuclear shuttling (**Fig. 5j-m**). Together these results not only indicate that PARPi-resistant TNBC display dependence on lysosomal pathways, but they also suggest a novel role for OA-NO_2_ in negative modulation of the lysosomal pathways.

## DISCUSSION

The unique phenotype of TNBC presents an unmet demand for improved targeted therapy. Despite being the most aggressive BC subtype that manifests poorer prognosis in patients, therapeutic strategies for TNBC have not evolved and consist of surgical removal of tumor(s), followed by adjuvant treatment with radiotherapy or chemotherapy which induces off-target organ damage, reduced bone density, and hair loss ^70–72^. The introduction of PARPi has revolutionized the treatment and increased survival rates for a small subset of TNBC patients presenting *BRCA* mutations ^6^.

As most patients develop resistance to PARPi ^7^, a better understanding of mechanisms that lead to PARPi resistance are desperately needed. The restoration of DDR pathways has been the predominantly studied mechanism underlying PARPi resistance ^7,37,73^. Importantly, alternative mechanisms of PARPi resistance are emerging that include autophagy as an important and druggable mechanism of PARPi resistance ^11,13,14^.

Autophagy-related PARPi resistance mechanisms are reported for ovarian and prostate cancers, but there is a significant gap in understanding whether autophagy is a PARPI resistance mechanism in TNBC ^13,74^. Herein, we address this gap in knowledge and present insights that are relevant to not only TNBC, but also multiple drug-resistant cancers. This study is unique in that it utilizes clinically relevant models of PARPi resistance in TNBC by generating novel *mutBRCA1* TNBC cell lines, since current studies of PARPi resistance in TNBC do not use patient-relevant *mutBRCA* TNBC cell lines ^11,12^. Notably, drug strategies targeting autophagy inhibition capitalize on recent observations that autophagy induction occurs in tumor cells in response to PARPi treatment ^8–13,16,38^, with autophagy inhibition resensitizing PARPi-resistant cells to PARP inhibition. Pharmacological and genetic inhibition of autophagy in concert with PARPi treatment can induce synergistic cytotoxicity in both *in vitro* and *in vivo* models ^10–13,74^. As a result, the clinical evaluation of autophagy as an exploitable therapeutic target has garnered increasing interest. The current data reports the mechanisms underlying how a safe orally bioavailable small molecule nitroalkene drug class can combat drug resistance driven by autophagy and the upregulation of the DDR ^75^.

Herein, we reinforce the importance of autophagy-lysosomal signaling in TNBC drug resistance. Like autophagy, lysosome-mediated chemoresistance is an understudied research area that is recently garnering clinical interest, highlighted by the recent development and testing of small-molecule lysosomal inhibitors ^76–78^. One notable transcription factor that modulates autophagy-lysosomal function is TFEB ^79^. Aside from regulating the expression of several autophagy proteins, such as p62 and LC3B, TFEB is also linked with increased lysosomal biogenesis and chemotherapeutic resistance in TNBC ^42,69^. The present report a) corroborates these previous findings of increased TFEB expression and a reliance on lysosomal biogenesis in drug-resistant TNBC cells, and b) demonstrates that TFEB is regulated by OA-NO_2_. These findings set the foundation for future investigation in lysosomal inhibition by OA-NO_2_ and other small molecule nitroalkenes.

The present data reveals that OA-NO_2_ inhibits the shuttling of TFEB to the nucleus. There is a correlation between levels of p62 and TFEB (**Supplemental Fig. 5e-g**), with knockdown of TFEB expression linked with increased p62 levels ^80–87^. It is possible that OA-NO_2_ can also impact intra-nuclear TFEB transcriptional regulatory functions, in addition to decreasing nuclear TFEB translocation, by alkylating the functionally-significant Cys105 and Cys 113 of p62. Given that TFEB nuclear localization is regulated by Cys212 oxidation ^88^, the direct regulation of TFEB by OA-NO_2_ alkylation of Cys212 might be an additional explanation for the present findings.

Aside from p62, additional autophagy-related signaling mediators have redox-sensitive cysteines, including ATG3, ATG7 ^89^, ATG4B ^90^, and TFEB ^88^, that would be sensitive to not only oxidation, but also alkylation and S-nitrosylation ^53,54^. Further investigation to determine the functional consequence of OA-NO_2_ targeting these proteins could be better defined by chemoproteomic-based screening strategies ^91,92^. This approach would also elucidate whether OA-NO_2_ targets other cysteine residues of p62, some of which have recently been described as susceptible to PTMs, including S-nitrosylation ^48–54^.

RNA-seq provides invaluable insight into potential mechanisms of PARPi resistance in TNBC. The present results align with previous reports, where olaparib-resistant cancer cell lines upregulate DNA damage responses and autophagy signaling autophagy pathways ^14,93^. These findings also align with prostate, ovarian, and breast cancer-related responses such as EMT and inflammatory upregulation in *in vitro* ^41,46,94^, *ex vivo* ^95^, and clinical scenarios ^96,97^. The clinical relevance of the PARPi-resistant cell lines reported herein is strengthened by previous identification of differences between PARPi-resistant *mutBRCA* samples ^98^ and specific responses to different PARPi ^99^. The current literature lacks perspective about genetic alterations in PARPi resistance in *mutBRCA* TNBC, both in patient samples and cell lines that develop acquired PARPi resistance. This study presents novel findings in not only two clinically relevant *mutBRCA* TNBC cells but also transcriptomic response differences for different PARPi. Further mining of the myriad of results coming from RNAseq analysis if different PARPi sensitive and resistant tumor cell populations will expand upon how one might better modulate PARPi resistance in TNBC.

Even though DNA-damaging agents such as PARPi induce autophagy, the relationship between genomic maintenance and autophagy is not entirely delineated ^100–102^. At the crossroads of genomic maintenance and autophagy, p62 plays a central role. As a multifaceted protein, p62 shuttles between the cytoplasm and nucleus, notably localizing to protein clusters in the nucleus ^103,104^. Dysfunctional autophagy leads to the accumulation of p62, which promotes mitochondrial damage, increased ROS production, and DNA damage ^105–107^. In addition, p62 impairs HR, promotes proteasomal degradation of filamin A, and decreases recruitment of Rad51 to DSBs ^102^.

Herein, we report evidence that affirms autophagy dependence as a PARPi resistance mechanism and present a novel, safe, clinical-stage therapeutic option for TNBC patients. The amplification of synergistic tumor cell killing by PARPi upon autophagy inhibition by OA-NO_2_ in PARPi-resistant TNBC has highlighted the clinical utility of targeting autophagy more effectively in not only chemotherapeutic-resistant TNBC but also in broader patient populations that could benefit from PARPi therapy, including those expressing HR proficiency or *wtBRCA1/2*. Compared to the risky autophagy inhibitors CQ and HCQ, OA-NO_2_ is an attractive candidate for chemotherapeutic applications, given its safety and favorable toxicology profiles.

## MATERIALS AND METHODS

### Cell Culture

HEK293T, HEK 293FT, MDA-MB-231, MDA-MB-436, and Hs578T cells (ATCC) were cultured at 37°C in 5% CO_2_ in DMEM (Gibco) supplemented with 10% FBS (Gibco), 100 units/mL penicillin/streptomycin (Gibco), nonessential amino acids (Gibco), and 2 mM GlutaMax (Gibco). p62 knockout (*p62^−/−^*) and wild-type (*p62^+/+^*) mouse embryonic fibroblasts (MEFs) were a kind gift from Eiji Warabi of the University of Tsukuba and Viktor Korolchuk of Newcastle University Institute. These MEFs were cultured in DMEM under the same conditions as described above. HCC1937 (ATCC) cells were cultured at 37 °C with 5% CO_2_ in RPMI-1640 (Gibco) supplemented with 10% FBS (Gibco), 100 units/ml penicillin/streptomycin (Gibco), nonessential amino acids (Gibco), and 2 mM GlutaMax (Gibco).

For p62 oligomer formation experiments, cells were maintained in serum starvation (SS) conditions at 0.5% DMSO for the duration of the treatment. For other experiments, cells were maintained in the same culture medium, except containing 5% FBS and 0.5% DMSO.

### Reagents and Antineoplastic Agents

Talazoparib (MedChemExpress), olaparib (Selleck Chemicals), niraparib (MedChemExpress), XRK3F2 (Selleck Chemicals), cycloheximide (CHX, Sigma Aldrich), and bafilomycin A1 (Cell Signaling Technology) were dissolved in dimethyl sulfoxide (DMSO). Fresh cis-Diammineplatinum (II) dichloride (Cisplatin) stocks were dissolved in sodium chloride solution (Gibco). CQ (Selleck Chemicals) and hydrogen peroxide (H_2_O_2_, 30% w/w, Sigma Aldrich) were dissolved in nuclease-free water suitable for cell culture. Oleic acid (OA) and OA-NO_2_ was synthesized as described previously ^108–111^. CP-8 was synthesized as previously described ^30^. (E)-10-nitro-octadec-9-en-17-ynoic acid (Alkynyl OA-NO_2_, alk-OA-NO_2_) was synthesized by ACME Bioscience/Frontage Lab. Pure OA, OA-NO_2_, alk-OA-NO_2_, and CP-8 were diluted in DMSO and added to cells after solvation in assay medium.

### Dose Response Assay

MDA-MB-231 (500 cells/well), MDA-MB-436 (500 cells/well), and HCC1937 (2000 cells/well) cells were plated in 96-well tissue culture plates one day prior to treatment (Day 1). Vehicle (0.5% DMSO in media) and treatment drugs were added to cells after solvation in the assay medium at the indicated concentrations. Drugs and vehicle were replenished three days later on Day 4. On Day 7, relative cell numbers were compared by measuring the luminescence signal generated by ATP using the CellTiter-Glo (Promega) assay per the manufacturer’s protocol. Luminescence values were calculated by the GloMax (Promega) microplate reader. Cell viability values were calculated by blank cell deductions and normalization to corresponding vehicle readings. For combination studies, synergy was calculated using SynergyFinder ^112^.

### Incucyte Cytotoxicity Assay

MDA-MB-231 cells were transduced with NucLight Red Lentivirus (Sartorius) and selected for cells expressing nuclear-restricted mKate2. After puromycin selection, cells were seeded at 500 cells/well into 96-well plates. After 24 hr, cells were treated with the indicated drugs and 250 nM Incucyte Cytotox Green (Sartorius) reagent to measure cell death. Three days later, the cells were replenished with media and reagents. The Incucyte Zoom (Sartorius) imaging system was used to record images every 4 hours. Live and dead cell quantification, red and green fluorescence respectively, was calculated using the manufacturer’s live cell analysis module.

### Immunoblotting

After treatments, cells were washed in ice-cold 1X phosphate buffered saline (PBS). Cell lysates were prepared by scraping on ice with cold RIPA Buffer with EDTA and EGTA (Boston Bioproducts) supplemented with Halt Protease and Phosphatase inhibitors (Thermo Fisher Scientific). Lysates were sonicated and subsequently centrifuged at 15,000 RPM for 10 min at 4°C. Protein concentrations were measured using the Pierce BCA protein assay kit (Thermo Fisher Scientific). Reducing 6X Laemmli sample buffer (Boston Bioproducts) was added to sample and incubated for 5 min at 95°C. An equal amount of protein lysates was loaded and separated on tris-glycine gels and transferred onto either polyvinylidene difluoride (PVDF) or nitrocellulose membranes. Membrane blocking was performed with either 5% bovine serum albumin (BSA) (Fisher Scientific) or 5% non-fat dry milk (Bio-Rad) in tris-buffered saline with 1% Tween 20 (1% TBST) for 1 hr at room temperature (RT). Primary antibodies (1:1000) were probed overnight in 5% BSA in TBST at 4°C. After 3 x 5 min washes in TBST, blots were incubated with secondary antibodies for 1 hr at RT. Blots were then washed 3 x 15 min prior to imaging. When probed with horseradish peroxidase (HRP)-conjugated secondary antibodies (1:2000, Rockland), blots were developed with SignalBright Max Chemiluminescent Substrate (Proteintech) using the Bio-Rad ChemiDoc imaging system. When probed with fluorescent secondary antibodies (1:10,000, LiCor), blots were developed using LiCOr Odyssey CLx Imaging system. Band quantifications were performed with ImageJ software.

For oligomer formation experiments, fresh 50mM N-ethylmaleimide (NEM) was added to lysis buffer in order to prevent new disulfide bond formation during lysis. Non-reducing 6X Laemmli sample buffer (Bioworld) was added to sample instead of reducing 6X Laemmli sample buffer. An equal amount of protein was run on 6-8% tris-glycine gels.

### Plasmids

pBABE-puro mCherry-EGFP-LC3B (Addgene #22418) was a kind gift from Jayanta Debnath and Prof. Roderick O’Sullivan ^43,113^. The plasmid was transformed into competent DH5α Max efficiency cells (Invitrogen) for plasmid amplification. Plasmid DNA was extracted and purified using ZymoPURE II Midiprep Plasmid Purification Kit (Zymo Research). The plasmid was transfected into MDA-MB-231 cells using the Fugene6 Transfection Reagent (Promega). mCherry-eGFP expressing cells were selected by puromycin dihydrochloride (Thermo Fisher Scientific). as well as by sorting with a BD FACSARIA at the Flow Cytometry Core at the Magee-Women’s Research Institute (Pittsburgh).

HA-p62 was a gift from Qing Zhong (Addgene plasmid # 28027) ^114^. The HA-p62 plasmid was transformed into competent DH5α Max efficiency cells (Invitrogen) for plasmid amplification. Plasmid DNA was extracted and purified using ZymoPURE™ II Midiprep Plasmid Purification Kit (Zymo Research). The plasmid was transfected into HEK293T cells using the Fugene6 Transfection Reagent (Promega). The following plasmids were a kind gift from Viktor Korolchuk: pLENTI6-V5-DEST empty vector, pLENTI6-V5-DEST Flag-p62, pLENTI6-V5-DEST Flag-p62 C105,113A ^48^. These plasmids were transformed into Stbl3 Chemically Competent cells (Invitrogen) for plasmid amplification. Plasmid DNA was extracted and purified using ZymoPURE™ II Midiprep Plasmid Purification Kit (Zymo Research).

### Animals

Animals used for this study were approved by and conducted according to the University of Pittsburgh Institutional Animal Care and Use Committee guidelines. MDA-MB-231 cells (1 × 10^6^) were injected into the mammary fat pad (left fourth gland) of 7-week-old female Nu/J mice in a volume of 20 μL of sterile saline as previously described ^29^. When tumors reached an average volume of 100 mm^3^, mice were randomized into groups and administered vehicle (5% DMSO, 30% PEG, 65% PBS), 0.33 mg/kg talazoparib, 15 mg/kg OA-NO_2_, or 0.33 mg/kg talazoparib in combination with 15 mg/kg OA-NO_2_ every day by gavage (200 μL) until experiment completion. The surgical procedure has been described previously ^115^.

### Proliferation Assay

An equal number of TNBC cells were seeded in multiwell plates and cultured for 4–5 days. Proliferation was assessed by fixing the plates for 30 min with formalin after which they were stained with 0.05% crystal violet. Wells were destained using 10% acetic acid. Absorbance (590 nm) was measured using a GloMax (Promega) microplate reader. Absorbance was normalized to vehicle wells.

### Cell Cycle Analysis

Unsynchronized TNBC cells were plated in 6-well plates for a final confluency of 70 - 90% by time of analysis. Cells were harvested by trypsinization and following inactivation of trypsin, cells were pelleted and washed with ice cold 1X PBS. Staining was done using the Propidium Iodide Flow Cytometry Kit (Abcam). Samples were fixed and stained following the manufacturer’s protocol. Samples were protected from light and placed on ice before direct analysis by flow cytometry. Data acquisition was performed on a Beckman Coulter CytoFlex 4L and data analysis was performed with FlowJo.

Unsynchronized TNBC cells were plated in 6-well plates for a final confluency of 70 - 90% by time of analysis. Cells were harvested by trypsinization and following inactivation of trypsin, cells were pelleted and washed with ice cold 1X PBS. Staining was done using the Propidium Iodide Flow Cytometry Kit (Abcam). Samples were fixed and stained following the manufacturer’s protocol. Samples were protected from light and placed on ice before direct analysis by flow cytometry. Data acquisition was performed on a Beckman Coulter CytoFlex 4L and data analysis was performed with FlowJo.

### RNA-Seq and Analysis

Total RNA was extracted from TNBC cells in nuclease-free water using RNeasy Plus Universal mini kit following manufacturer’s instructions (Qiagen). RNA concentration and quality was assessed via a NanoDrop spectrophotometer (Thermo Fisher Scientific). Samples were then packaged and shipped to Novogene for further processing and sequencing.

### Library construction and sequencing

Messenger RNA (mRNA) was purified from total RNA using poly-T oligo-attached magnetic beads. After fragmentation, the first strand cDNA was synthesized using random hexamer primers, followed by the second strand cDNA synthesis using either dUTP for directional library or dTTP for non-directional library. The library was checked with Qubit (Thermo Fisher Scientific) and real-time PCR for quantification and bioanalyzer for size distribution detection respectively. Quantified libraries were pooled and sequenced on Illumina (Illumina Inc.) platforms, according to effective library concentration and data amount. Clustering of the index-coded samples was performed according to the manufacturer’s instructions. After cluster generation, the library preparations were sequenced on an Illumina platform and paired-end reads were generated.

### Quality control (QC), read mapping, and quantification of gene expression level

First, raw data (raw reads) of fastq format were processed through fastp software. Clean data (clean reads) were obtained by removing reads containing adapter, reads containing ploy-N and low-quality reads from raw data. Concurrently, Q20, Q30 and GC content (clean data) were calculated. All downstream analyses were based on high quality clean data. Index of the reference genome was built and paired-end clean reads were aligned to the reference genome using Hisat2 v2.0.5 ^116^. Quantification of the gene expression level was done via featureCounts v1.5.0-p3 ^117^. The Fragments Per Kilobase of transcript sequence per Millions base pairs sequenced (FPKM) of each gene was calculated based on the length of and the reads count mapped to the gene.

### Principle component, clustering, and pathway analyses

To identify differently expressed genes and calculate fold changes between groups, we used DESeq2 ver.1.44.0 ^118^. Significantly differentially expressed genes (DEGs) were defined as genes with false discovery rate (FDR) q-value of less than .05. To explore the expression clusters of the PARPi-resistant mutBRCA cells, we performed unsupervised hierarchical clustering analysis and Principal Component Analysis (PCA) with the first two components. TPM log2 was used to observe unbiased hierarchical clustering using R package ComplexHeatmap ver.2.20.0 ^119^. PCA was performed by building the DESeq2 model and the variance stabilizing transformation (VST). Venn diagram plots were generated using the VennDiagram package ver.1.7.3 ^120^. In addition, we performed gene set enrichment analysis (GSEA) ^121^ or Weighted Concept Signature Enrichment Analysis (WCSEA) with the IndepthPathway R package ver.1.0 ^122^ to identify the signaling pathways characteristic of the cells in different PARPi models. For the gene signature data, we used the hallmark or canonical pathways (C2CP) downloaded from Molecular Signature DataBase (MSigDB) ^123^. Otherwise, we generated our own data with gene signatures related to autophagy, DNA Repair, and lysosome. We calculated the mean of normalized enrichment score (NES) and FDR from the pairwise GSEA or WCESA and set the FDR q-value to 0.05 (5%) as the threshold to identify significantly enriched pathways. Data visualization was performed using the ggplot2 package ver.3.5.1 ^124^.

### Real Time-qPCR

RT-qPCR was performed as described previously ^125^. RNA was isolated using a Nucleospin RNA isolation kit (Takara Bio USA) and RNA concentration and quality assessed via a NanoDrop spectrophotometer (Thermo Fisher Scientific). cDNA was synthesized with the SuperScript III First-Strand Synthesis System for RT-PCR (Invitrogen) using random hexamers per the manufacturer’s protocol. SYBR green-based RT-PCR was performed using the CFX384 Real-Time qPCR Detection System (Bio-Rad) and respective primers. The comparative Ct method was used for data analysis with GAPDH as the comparator gene.

### Irradiation

Experiments were conducted on cells dosed with 0 or 5 Gy utilizing a benchtop CellRad+ x-ray ɣ-Irradiator (Precision X-Ray) with a dose rate of 1.3 Gy/min.

### Click Chemistry

After 1 hr treatment of alkynyl probe, cells were trypsinized and washed with 1X cold PBS. Samples were then gently resuspended in 100 μL lysis buffer (100 mM Sodium phosphate, 50 mM HEPES, pH 7.4, 1% Triton X-100) supplemented with Halt Protease and Phosphatase inhibitors (Thermo Fisher Scientific) and incubated on ice for 15-30 min. Lysates were sonicated and subsequently centrifuged at 15,000 RPM for 10 min at 4°C. Supernatant were then transferred to fresh microcentrifuge tubes and diluted up to 460 μL with HEPES buffer without Triton (100 mM Sodium phosphate, 50 mM HEPES, pH 7.4). For the click chemistry reaction, the following chemicals were added to the lysate in order: biotin azide (0.02 mM final concentration, Vector Laboratories), tris(benzyltriazolylmethyl)amine (2 mM final concentration, TBTA, Vector Laboratories), copper (II) sulfate (1 mM final concentration, Sigma Aldrich), and sodium ascorbate (10 mM final concentration, Sigma Aldrich). The samples were vortexed briefly after the addition of each chemical, with a final sample volume of 500 μL. Samples were incubated for 1 hr at RT while rocking and protected from light. 8 μL of 0.5 M EDTA was then added to stop the click reaction and the samples were acetone precipitated. Protein pellets were allowed to air-dry on ice, resuspended in 200 μL of 6 M urea + 1% SDS by sonication, and then diluted up to 400 μL in 1X PBS. Magnetic streptavidin beads (Thermo Fisher Scientific) were washed according to the manufacturer’s protocol, and 100 μL of the beads were added per sample. The samples were incubated with the beads for 1 hr at RT by rocking. The beads were sequentially washed with 1% SDS in PBST (5 × 5 min) with rocking, allowing samples to attach to the magnetic rack between each wash. For immunoblot analysis, 1X reducing Laemmli sample buffer (Boston Bioproducts) diluted in lysis buffer was added to the beads and incubated at 95°C for 5 min. After samples were attached to the magnetic rack, the lysate was eluted from the beads and transferred to a fresh tube for further analysis.

### Lentiviral Transduction

Stable expression of FLAG-p62 and FLAG-Cys105A-C113Ala p62 in *p62^−/−^*MEFs was achieved through lentiviral transduction. Semi-confluent HEK293T cells were co-transfected with either empty or p62 lentiviral expression vectors and 3rd generation packaging system plasmids (Invitrogen). Following seeding in 6-well plates 24 hr prior, 200 µL of virus was added to the cells with polybrene (6 μg/ml, Millipore Sigma) and incubated for 24 hr. Drug selection with blasticidin (4 μg/ml, Invitrogen) was performed 3 days after transduction. After confirmation of p62 expression, transduced MEFs were then maintained in lower levels of until seeding for experimental purposes as described previously ^48^.

SQSTM1/p62 shRNA were received from the TRC library (Millipore Sigma), with the following targeting sequences: TRCN0000007235 (sh#1): GCAGATGAGAAAGATCGCCTT; TRCN0000007236 (sh#2): CCGAATCTACATTAAAGAGAA; TRCN0000007237 (sh#3): CCTCTGGGCATTGAAGTTGAT. shRNA was transformed into DH5α Max efficiency cells (Invitrogen). pLKO.1 scramble (Scr) shRNA was used as controls. The expression or shRNA plasmids were co-transfected with the virus package plasmids, psPAX2 (Addgene, #12260) and pMD2.G (Addgene, #12259), to HEK293FT cells for virus production. Cells were seeded in 6-well plates the day before virus transduction. 200 µL of virus was added to the cells with polybrene (0.8 µg/mL, Millipore Sigma) and incubated for 24 hr. Drug selection with puromycin (2 μg/ml Gibco) was performed 3 days after transduction.

### Cycloheximide degradation assay

*p62^−/−^* MEFs stably expressing FLAG-tagged wild type or Cys105A.C113Ala p62 were seeded in 6-well plates until 70 to 80% confluent. Cells were pre-treated with H_2_O_2_ (200 µM) and either CQ (20 µM) or OA-NO_2_ (5 µM) in serum-free media. 1 hr later, cells were given vehicle (0.5% DMSO) or CHX (50 μg/ml) treatment in serum-free media. Cell lysate was then collected at 0, 2, 4 and 6 hr timepoints for analysis of p62 degradation as described previously ^48^.

### LysoTracker Green DND-26 Flow Cytometry Assay

Quantification of lysosomal mass was performed similarly as described previously ^69^. Semi-confluent TNBC cells in 6-well plates were trypsinized and washed with 1X PBS. Cells were subsequently stained with 50 nM LysoTracker Green DND-26 (Thermo Fisher Scientific), rocking gently for 15 min at 37°C. Cells then were washed and resuspended in cold PBS supplemented with 2% fetal bovine serum. Samples were covered and placed on ice before direct analysis by flow cytometry. Data acquisition was performed on a Beckman Coulter CytoFlex 4L and data analysis was done with FlowJo.

### Nuclear fractionation

Extraction of the nuclear cell fraction was done using the NE-PER Nuclear and Cytoplasmic Extraction Reagents (Thermo Fisher Scientific) according to the manufacturer’s instructions. Following extraction of nuclear and cytoplasmic fractions, lysates were prepared for immunoblotting as described above.

### Statistical Analysis

GraphPad Prism (version 10) was used to perform statistical analysis. Unless otherwise noted, data represent the mean ± SEM from three or more independent experiments. IC50 values were calculated using a non-linear dose response variable slope mode. One-way ANOVA tested significance for multiple groups with Tukey posttest for multiple comparisons between groups or by Student’s unpaired t-test when groups were less than three. Two-way ANOVA was used to test for significance between multiple groups when indicated. Differences were considered significant at #p < 0.1, *p < 0.05, **p < 0.01, ***p < 0.001, ****p < 0.0001).

---

**Supplemental Figure 1:**
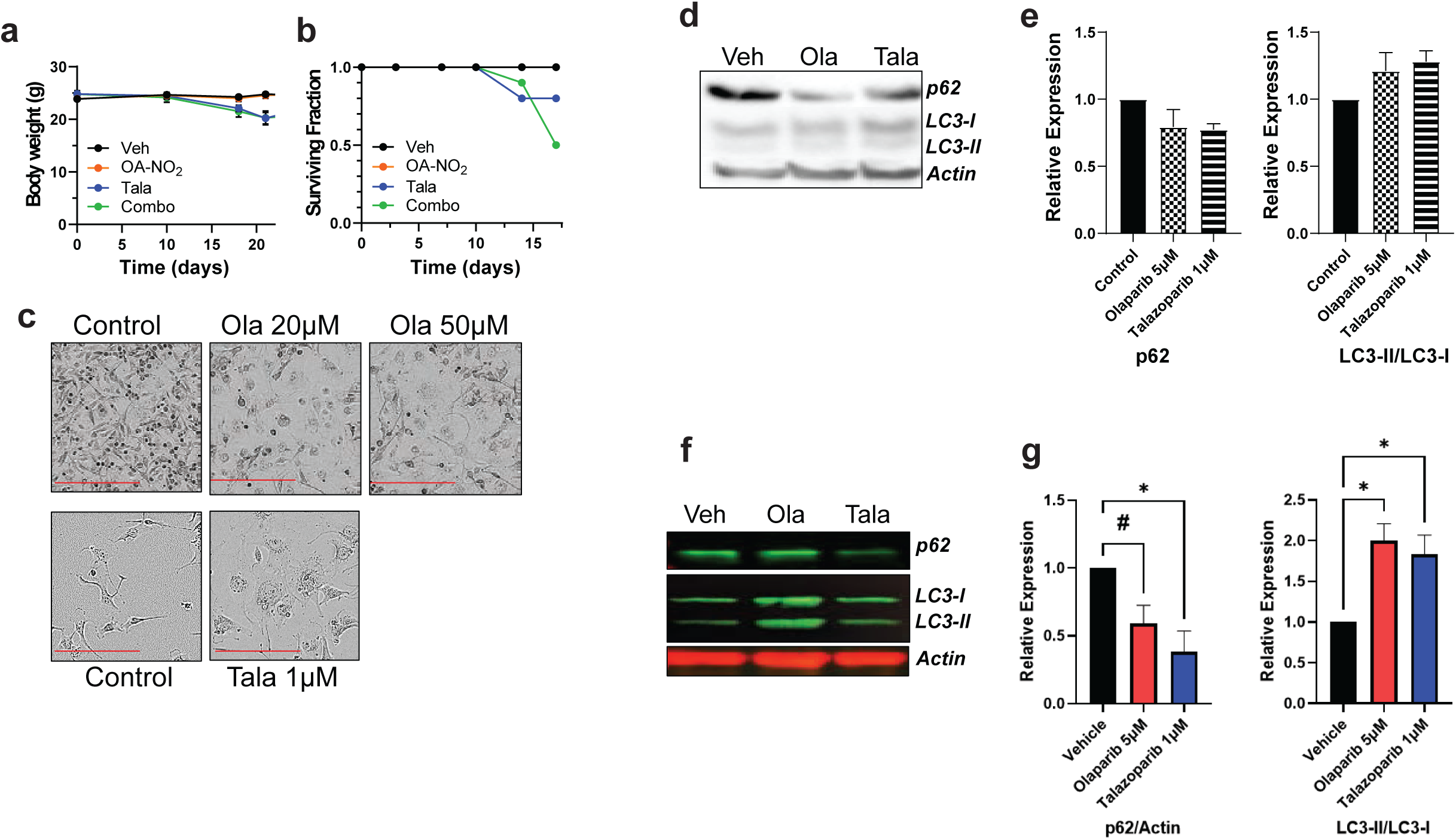
PARPi induce hypertrophy-associated autophagy

**Supplemental Figure 2:**
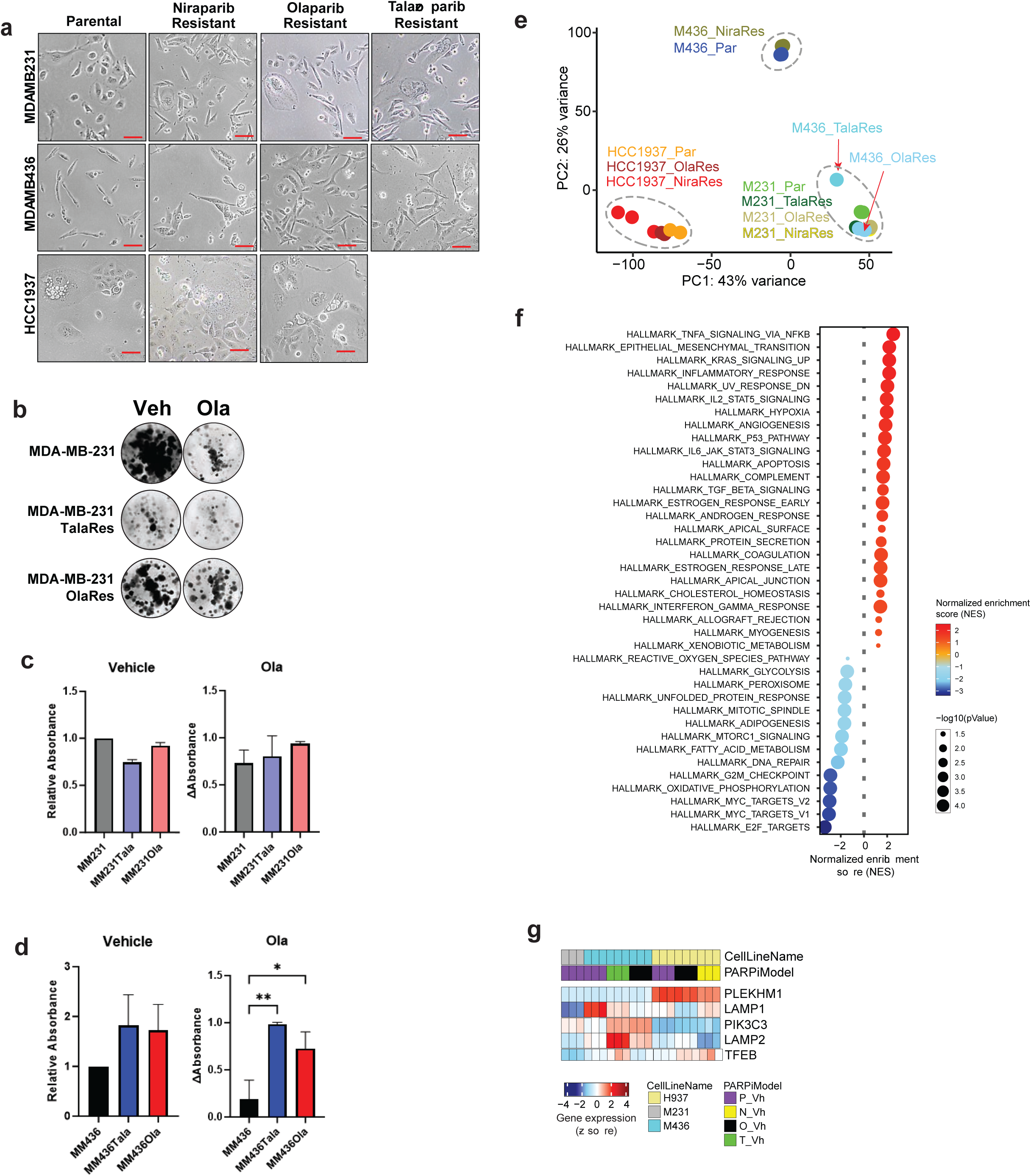
PARPi-resistant TNBC Present Morphological and Genetic Differences from Parental Cells.

**Supplemental Figure 3:**
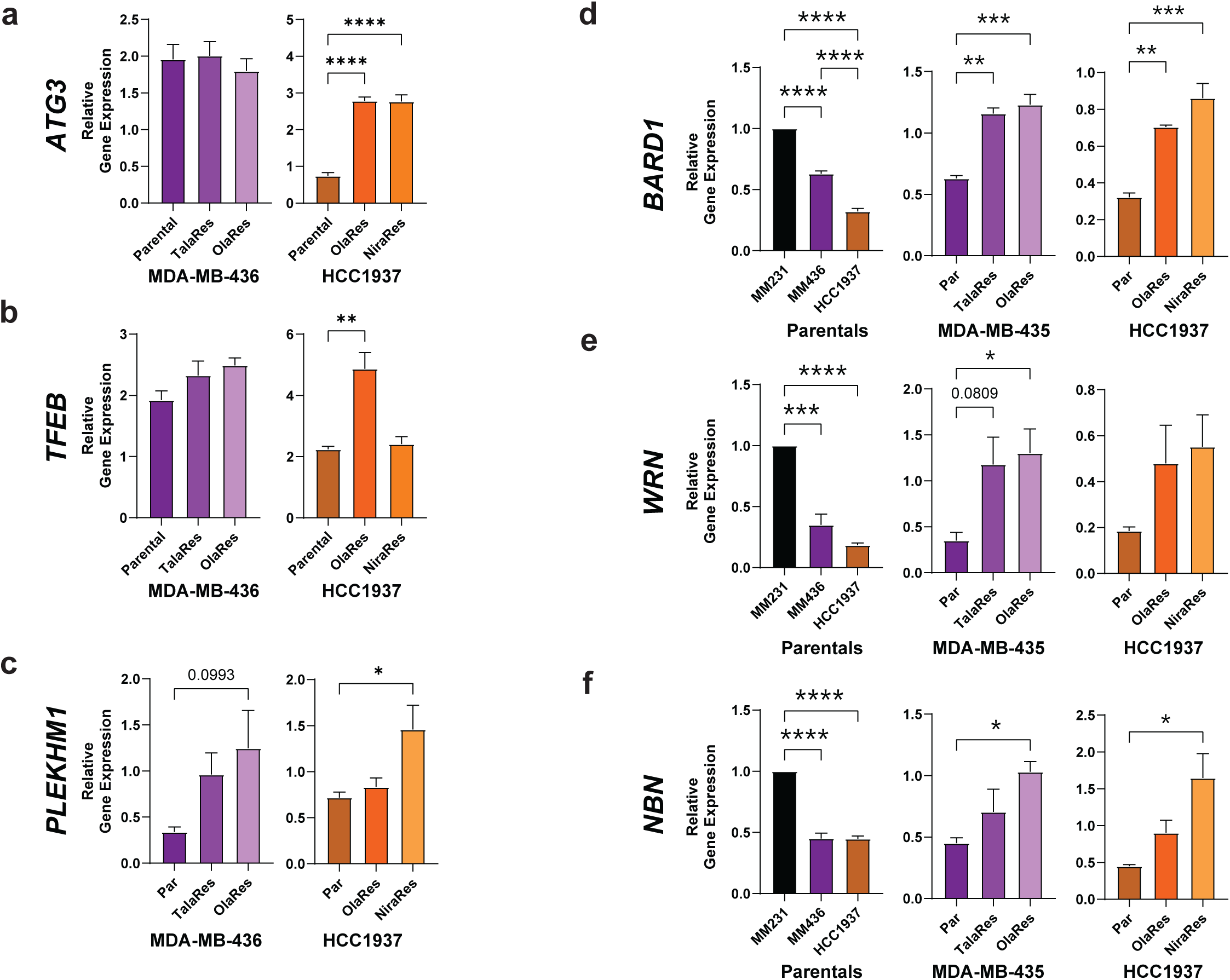
PARPi-resistant TNBC Present Differentially Expressed Genes.

**Supplemental Figure 4:**
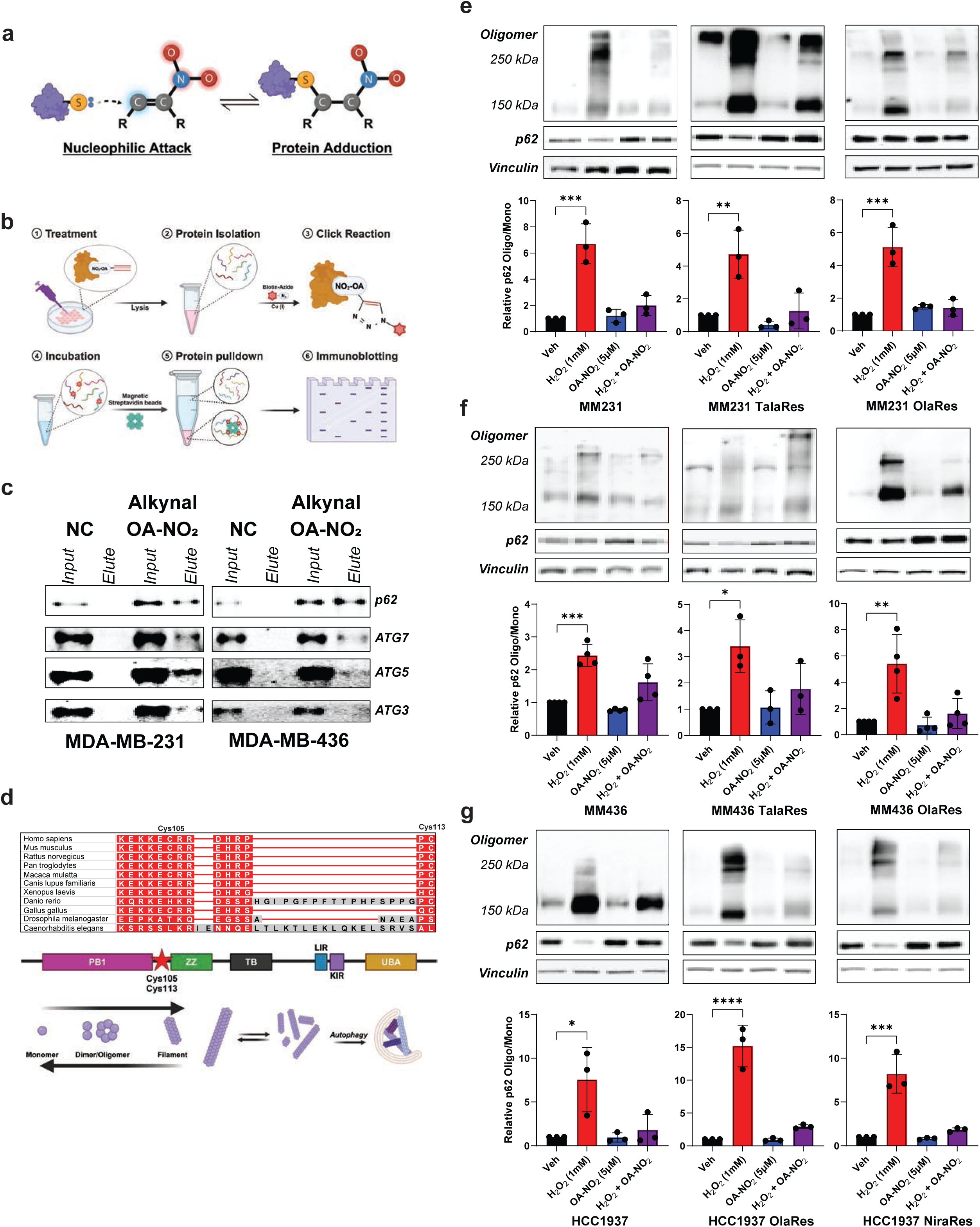
O A-NO_2_ Targets p6 2 in Parental and PARPi-Resistant TNBC.

**Supplemental Figure 5:**
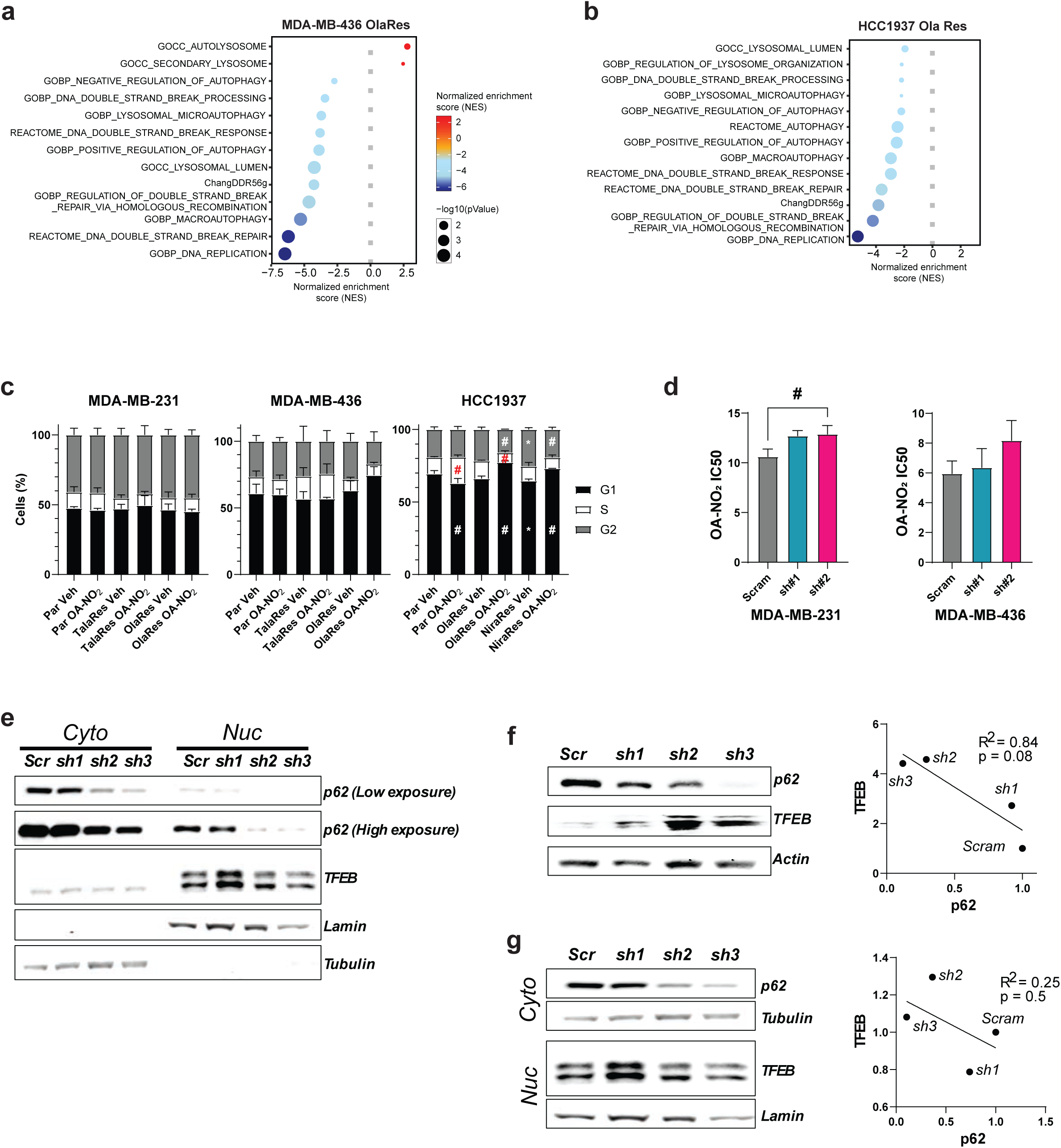
Knockdown of OA-NO2 Target p62 Correlates to Increased TFEB Expression.

**Supplemental Figure 6:**
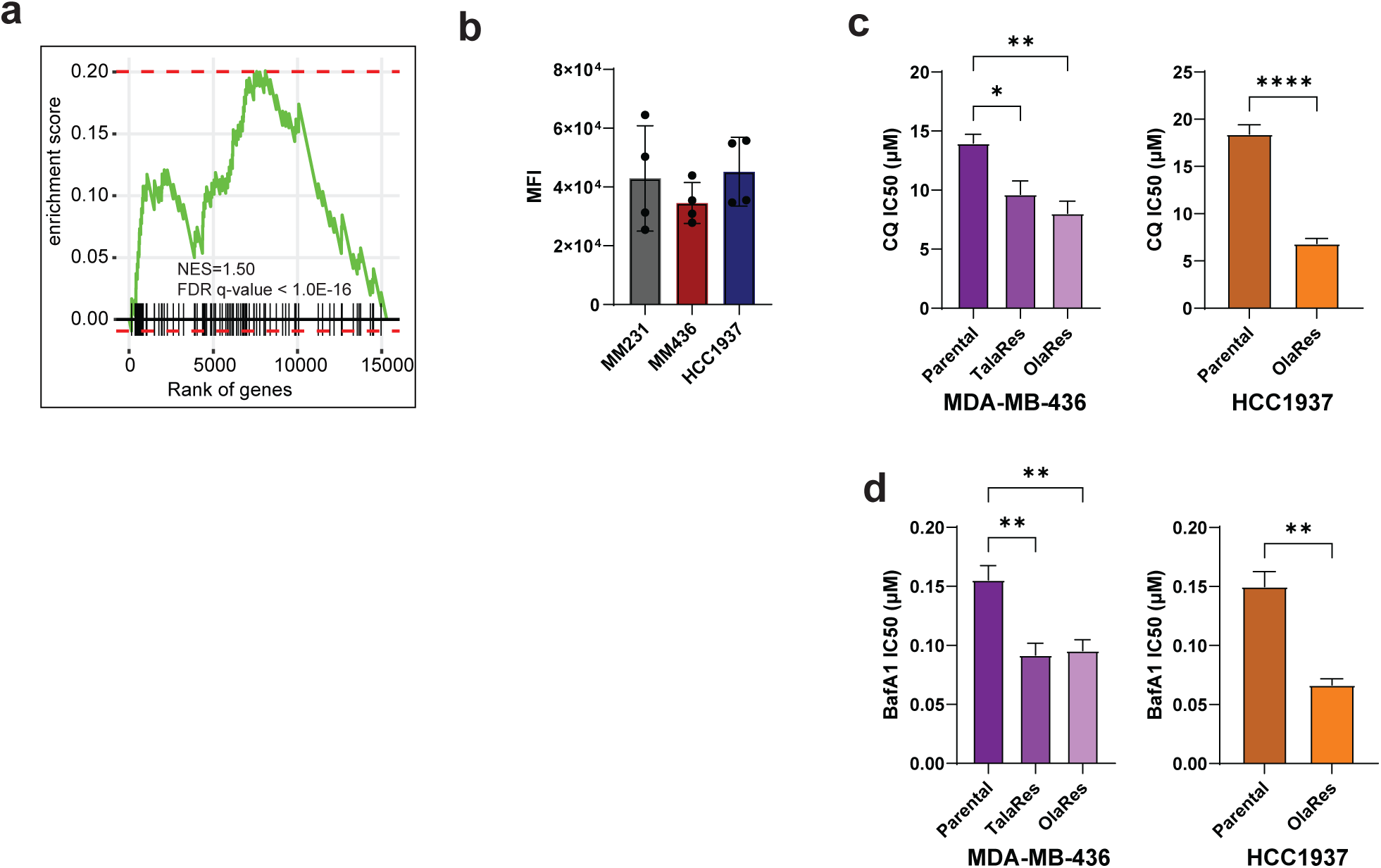
Increased Lysosome Dependence in PARPi-Resistant TNBC.

